# *Staphylococcus aureus* Delta Toxin Modulates both Extracellular Membrane Vesicle Biogenesis and Amyloid Formation

**DOI:** 10.1101/2023.03.23.533957

**Authors:** Xiaogang Wang, Divakara SSM Uppu, Seth W. Dickey, Dylan J. Burgin, Michael Otto, Jean C. Lee

**Affiliations:** Division of Infectious Diseases, Department of Medicine, Brigham and Women’s Hospital and Harvard Medical School, 181 Longwood Avenue, Boston, MA USA 02115; Pathogen Molecular Genetics Section, Laboratory of Bacteriology, National Institute of Allergy and Infectious Diseases, National Institutes of Health, Bethesda, MD, USA 20814; Department of Veterinary Medicine, Virginia-Maryland Regional College of Veterinary Medicine, University of Maryland, College Park, MD, USA 20740

**Keywords:** *Staphylococcus aureus*, extracellular membrane vesicles, phenol-soluble modulins, delta-toxin, amyloid fibrils

## Abstract

*Staphylococcus aureus* secretes phenol-soluble modulins (PSMs), a family of small, amphipathic, secreted peptides with multiple biologic activities. Community-acquired *S. aureus* strains produce high levels of PSMs in planktonic cultures, and PSM alpha peptides have been shown to augment the release of extracellular membrane vesicles (MVs). We observed that amyloids, aggregates of proteins characterized by a fibrillar morphology and stained with specific dyes, co-purified with MVs harvested from cell-free culture supernatants of community-acquired *S. aureus* strains. δ-toxin was a major component of amyloid fibrils that co-purified with strain LAC MVs, and δ-toxin promoted the production of MVs and amyloid fibrils in a dose-dependent manner. To determine whether MVs and amyloid fibrils were generated under in vivo conditions, we inoculated mice with *S. aureus* harvested from planktonic cultures. Bacterial MVs could be isolated and purified from lavage fluids recovered from infected animals. Although δ-toxin was the most abundant PSM in lavage fluids, amyloid fibrils could not be detected in these samples. Our findings expand our understanding of amyloid fibril formation in *S. aureus* cultures, reveal important roles of δ-toxin in amyloid fibril formation and MV biogenesis, and demonstrate that MVs are generated in vivo in a staphylococcal infection model.

**Importance:** Extracellular membrane vesicles (MVs) produced by *Staphylococcus aureus* in planktonic cultures encapsulate a diverse cargo of bacterial proteins, nucleic acids, and glycopolymers that are protected from destruction by external factors. δ-toxin, a member of the phenol soluble modulin family, was shown to be critical for MV biogenesis. Amyloid fibrils co-purified with MVs generated by virulent, community-acquired *S. aureus* strains, and fibril formation was dependent on expression of the *S. aureus* δ-toxin gene (*hld*). Mass spectrometry data confirmed that the amyloid fibrils were comprised of δ-toxin. Although *S. aureus* MVs were produced in vivo in a localized murine infection model, amyloid fibrils were not observed in the in vivo setting. Our findings provide critical insights into staphylococcal factors involved in MV biogenesis and amyloid formation.

## Introduction

*Staphylococcus aureus* is a primary cause of invasive infections in humans, such as bacteremia, endocarditis, pneumonia, and surgical wound infections, leading to morbidity, mortality, and excessive healthcare costs (1). By employing a diverse array of surface-associated and secreted virulence factors, *S. aureus* colonizes and invades host tissues and evades the host immune response (2–7). Lacking secretion systems (T3SS, T4SS, and T6SS) that transport Gram-negative bacterial virulence factors directly into host cells (8), *S. aureus* secretes its exoproteins to the external environment (8–11), where they may be inactivated by neutralizing antibodies or enzymes with proteolytic or hydrolytic activities. However, *S. aureus* also generates extracellular membrane vesicles (MVs), and virulence determinants packaged within MVs (12–17) are protected from destruction by external factors.

MVs are spherical membrane nanoparticles that are released by prokaryotes, eukaryotes, and archaea (18). The cargo of *S. aureus* MVs includes cytosolic, surface, and membrane proteins, as well as nucleic acids, glycopolymers, and secreted proteins, such as pore-forming toxins, superantigens, and proteases (12–17). We and others have shown that purified MVs are cytolytic to multiple cell types (12, 14, 15, 19), induce atopic dermatitis-like inflammation in mice (19, 20), and elicit the production of pro-inflammatory mediators (16, 19–23). MV production is enhanced by environmental stresses typically encountered by *S. aureus* during infection (24–26). Although many *S. aureus* strains produce MVs in vitro (12–15, 17, 23, 27), the generation of MVs in vivo during staphylococcal infection has not yet been documented.

Although the biogenesis of MVs generated by Gram positive bacteria remains poorly understood, phenol-soluble modulins (PSMs) have been shown to be critical for MV release in *S. aureus*. PSMs, produced by multiple staphylococcal species, are genome-encoded, amphipathic peptides with alpha-helical structures (28, 29). They are classified into two types: the α-type peptides with 21 to 26 amino acids (PSMα1, α2, α3, α4, and δ-toxin) and the β-type peptides with 44 amino acids (PSMβ1 and β2) (30). PSMα1-4 peptides enhance the release of MVs by altering the cell membrane due to their surfactant-like characteristics, thus increasing membrane fluidity and promoting MV biogenesis (14, 31). Synthetic PSMα3 at concentrations ≥25 μg/ml lysed MVs purified from *S. aureus* strain LAC (31). Under appropriate conditions, various PSM peptides have been shown to form amyloid fibrils alone or in combination (32–36). Most of the latter studies were performed in vitro with synthetic PSM peptides or under biofilm growth conditions.

In this report, we investigated the roles of the PSMα peptides, PSMβ peptides, and δ-toxin in the generation of *S. aureus* MVs and evaluated the relationship between PSM production and amyloid formation under planktonic culture conditions. Both PSMα peptides and δ-toxin enhanced *S. aureus* MV production, and δ-toxin formed amyloid fibrils that co-purified with MVs harvested from culture supernatants. MVs were produced in vivo in an *S. aureus* air pouch infection model, but amyloid fibrils were not detected in lavage fluids, suggesting that amyloid formation did not occur in vivo. Our study provides fundamental insights into the role of *S. aureus* PSM peptides in MV biogenesis and amyloid fibril formation in planktonic cultures.

## Results

### Strain-dependent production of MVs and amyloid fibrils in planktonic *S. aureus* cultures

We purified MVs from culture supernatants of hospital-acquired *S. aureus* isolates (ST5 strain N315, ST36 strain Sanger 252, and ST30 strain MN8), as well as the more virulent community-acquired *S. aureus* isolates (ST8 strain LAC, ST1 strain MW2, and ST59 strain NRS483). Negatively stained samples imaged by transmission electron microscopy (TEM) revealed diverse MV morphologies (**Fig. 1**). MVs generated by strains MN8, Sanger 252, and N315 varied somewhat in their appearance, consistent with previous reports (37). However, MV preparations purified from community-acquired *S. aureus* strains MW2, ST59, and LAC were distinctive as they contained an abundance of amyloid-like fibrils (**Fig. 1**).

**Figure 1.**
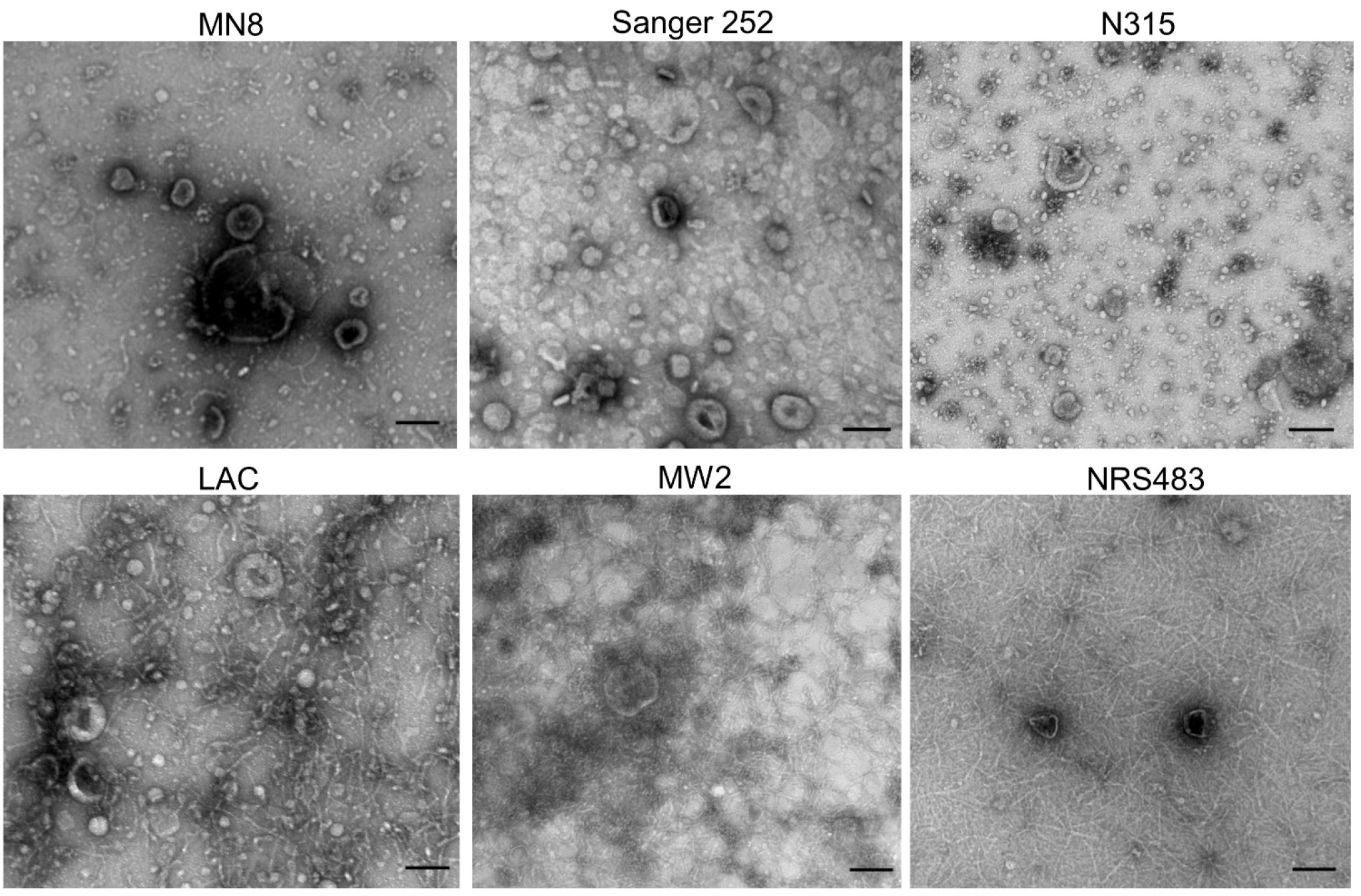
Electron micrographs of MVs purified from the post-exponential cultures of indicated *S. aureus* strains. MN8, Sanger 252, and N315 are hospital-acquired *S. aureus* strains, whereas strains LAC, MW2, and NRS483 are community-acquired isolates. Scale bar, 100nm. Images are representative of three independent experiments.

Despite employing a combination of tangential flow filtration, ultracentrifugation, density-gradient ultracentrifugation, and diafiltration to achieve MV purification, fibrils co-purified with MV preparations from the prototype *S. aureus* USA300 strain LAC. We considered that fibrils visualized by TEM could be an artifact of our MV purification process. To address this, we prepared additional samples wherein we omitted the step of concentrating the filter-sterilized culture supernatants by tangential flow filtration, and we did not purify the MVs via Opti-prep gradient ultracentrifugation and diafiltration. As shown in **Fig. S1A and 1B**, abundant fibrils were observed by TEM in the crude LAC MV pellet, but not in the crude Sanger 252 sample. To determine whether the observed fibrils were amyloid in nature, purified MV samples from each strain were stained with the amyloid-specific dye thioflavin T (ThT) (38). MVs generated by strain LAC displayed a dose-dependent increase in fluorescence that was lacking in MVs prepared from strain Sanger 252 (**Fig. S1C**). These data demonstrate that amyloid fibrils were formed in planktonic cultures of community-acquired *S. aureus* strains, and that the fibrils co-purified with MVs generated in vitro.

### δ-toxin plays a dominant role in amyloid formation in *S. aureus* planktonic cultures

Because previous studies demonstrated that either synthetic PSMs or PSMs produced in *S. aureus* biofilms formed amyloid fibrils in vitro (32–36, 39, 40), we hypothesized that PSMs contributed to amyloid fibril formation in planktonic cultures of *S. aureus* LAC. The expression of PSMs is growth phase dependent, and *agr* dysfunctional mutants lack detectable PSM production (28). Accordingly, MVs harvested from exponential phase cultures of strain LAC were not associated with fibrils (**Fig. S2A**). Likewise, MVs purified from post-exponential cultures of JE2Δ*agr* (**Fig. S2B**) were free of fibrils. In contrast, MV samples purified from post-exponential cultures of the pore-forming toxin mutant LACΔ*lukAB*Δ*hlgACB*Δ*lukED*Δ*pvl*Δ*hla* (41) (**Fig. S2C**) or JE2Δ*atl* (**Fig. S2D**) showed abundant fibrils, indicating that neither cytolysins nor cytoplasmic proteins released by the major autolysin Atl modulate fibril formation (42).

To investigate whether PSMs play a role in amyloid fibril formation, we purified MVs from culture supernatants of the wild-type (WT) strain LAC and LAC PSM mutants. Fibrils were present in MV samples purified from post-exponential cultures of LAC, LACΔ*psmα*, LACΔ*psmβ*, and LACΔ*psmα/β* **(Fig. 2A)**. In contrast, fibrils were absent from samples lacking δ-toxin (LACΔ*hld* and LACΔ*psmα/β*Δ*hld*). To rule out the possibility that fibrils might be removed during the various MV purification steps, crude MVs were pelleted by ultracentrifugation from filter-sterilized culture supernatants, suspended in PBS, and imaged by TEM. Consistently, fibrils were observed in the crude samples prepared from WT LAC, LACΔ*psmα*, LACΔ*psmβ*, and LACΔ*psmα/β*, but fibrils were absent in the *LAC*Δ*hld* samples (**Fig. S3A**). Likewise, we evaluated fibril formation from LAC and its mutants grown to stationary phase (8 h), and similar results were observed (**Fig. S3B**). PSM concentrations in culture supernatants of WT strain LAC grown to exponential (4 h), post-exponential (4.5 h), or stationary (8 h) growth phase are shown in **Fig. 2B**. At each timepoint, δ-toxin was the most abundant PSM peptide detected.

**Figure 2.**
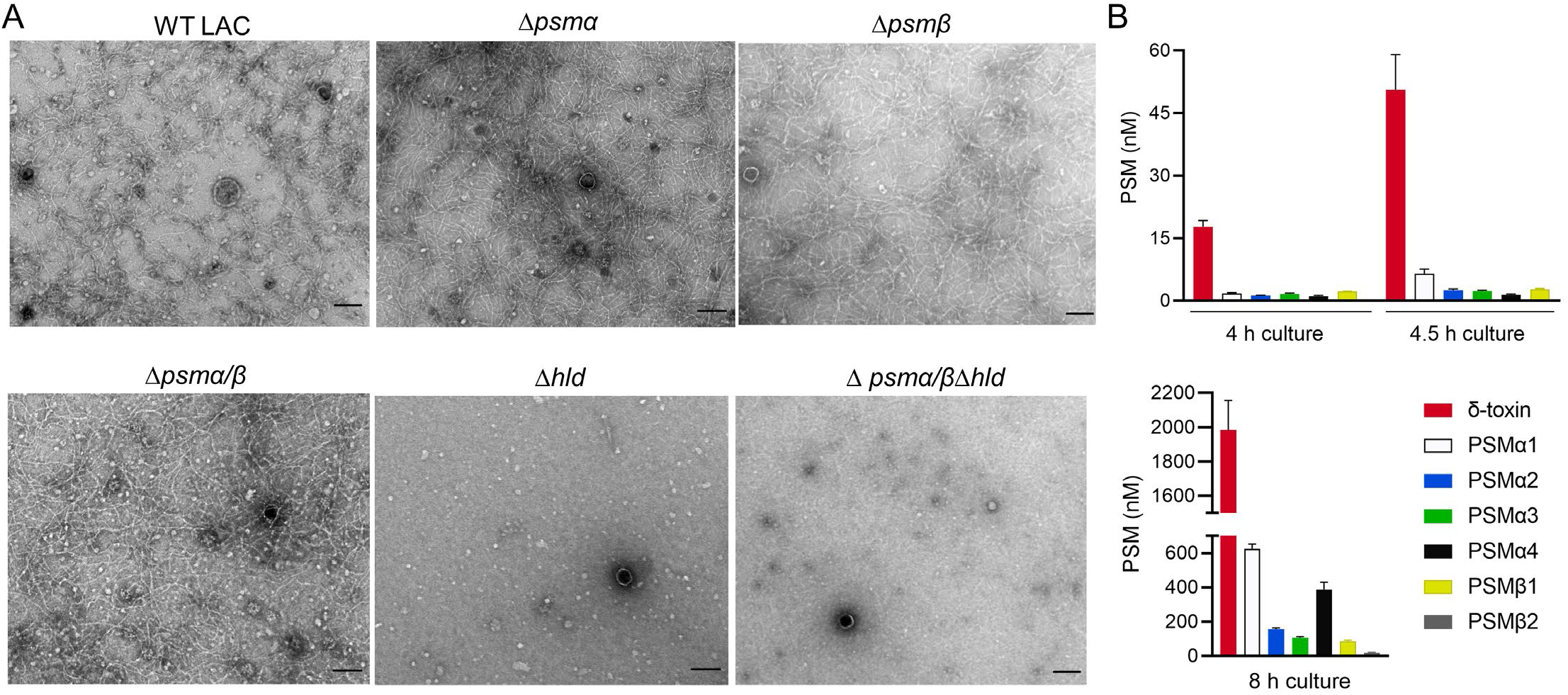
Effects of PSMs on amyloid fibril formation in post-exponential *S. aureus* LAC cultures. (A) MVs purified from WT LAC or indicated *psm* mutants were negatively stained and imaged by TEM. The experiments were performed at least twice with different batches of MV samples, and a representative image for each sample is shown. Scale bar, 100 nm. (B) The concentrations of individual PSM peptides in supernatants collected from exponential (4 h), post-exponential (4.5 h), or stationary phase (8 h) cultures of strain LAC. Values shown are means + SEM.

We complemented *LAC*Δ*hld* with pTX*-hld* (43), which allowed us to control the expression of *hld* by induction with 0, 0.1, 0.3, or 0.5% xylose. When the cultures reached the post-exponential growth phase, the MVs from each culture were purified and imaged by TEM. Fibrils were observed in MV samples purified from cultures with *hld* induction by xylose concentrations ≥0.3% (**Fig. 3A**).

**Figure 3.**
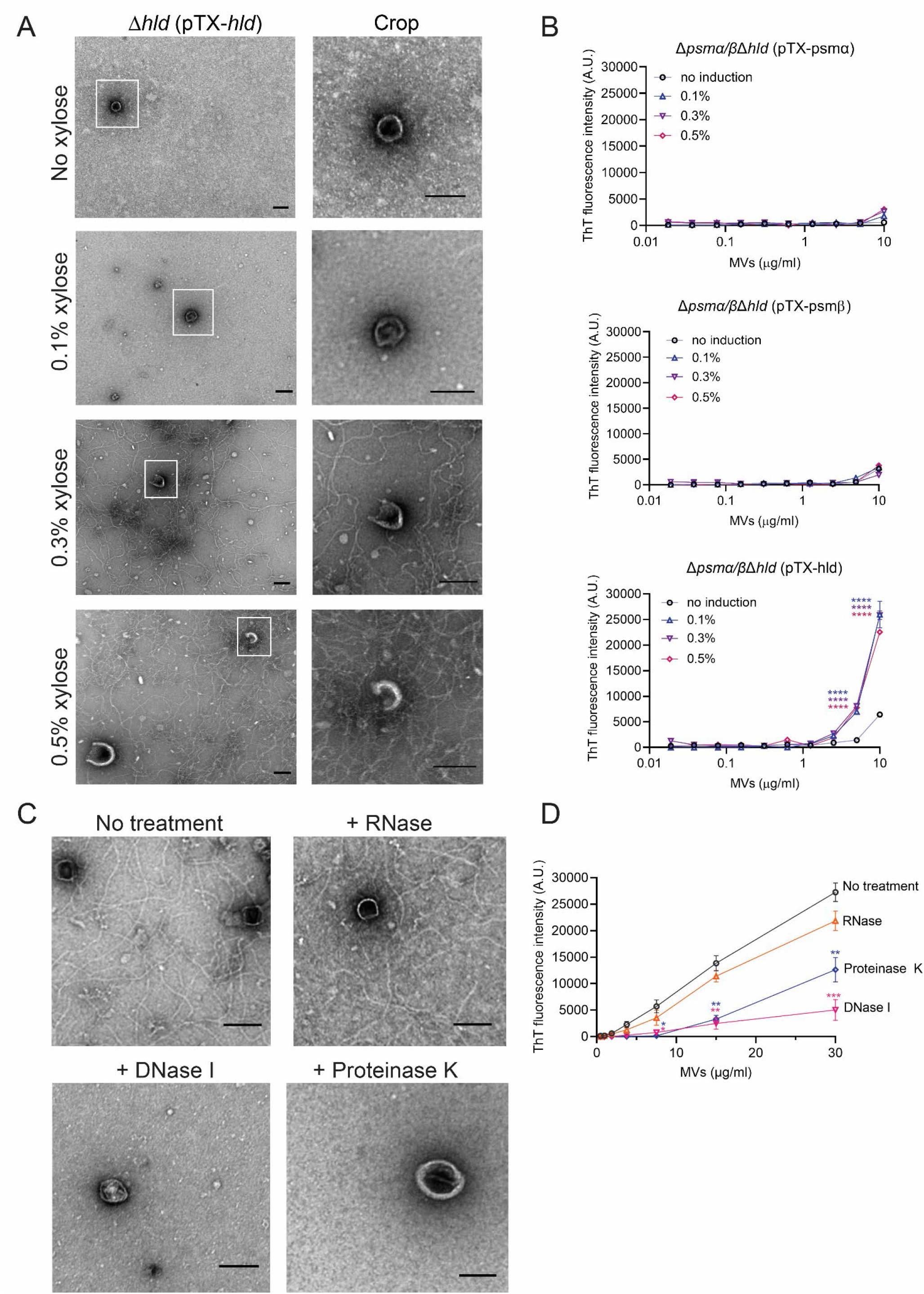
The formation of amyloid fibrils in post-exponential cultures of *psm* mutant strains complemented with *psmα, psmβ*, or *hld*. (A) Electron micrographs of MV samples purified from cultures of the LACΔ*hld* mutant expressing *hld* under the control of a xylose-inducible promoter. A cropped and magnified field from each image is shown. (B) ThT fluorescence analyses of MV samples purified from the Δ*psmα/βΔhld* mutant that was complemented with various *psm* genes and induced with increasing concentrations of xylose. (C) Electron micrographs of MV samples purified from *S. aureus* LAC cultivated in the presence or absence of 200 μg/ml DNase I, RNase, or proteinase K. MVs from the samples were negatively stained and imaged by TEM. Scale bars, 100 nm. (D) Fluorescence of ThT-stained MV samples purified from treated or untreated LAC cultures. ThT fluorescence was expressed as the mean ± SEM (n=3). Data were compared to the uninduced or untreated sample by the Student *t*-test. test. \*\**P* <0.01, \*\*\**P* < 0.001, \*\*\*\**P* < 0.0001.

To confirm that δ-toxin was solely responsible for amyloid fibril formation, the *LAC*Δ*psm*α/*β*Δ*hld* mutant was complemented with *psmα1-4, psmβ1-2*, or *hld* using inducible vector pTX (43). MVs from each culture were stained with ThT. Samples from the Δ*psm*α/*β*Δ*hld* (pTX-*hld*) strain induced with 0.1, 0.3, or 0.5% xylose exhibited significant fluorescence at MV concentrations ≥5 μg/ml (**Fig. 3B**). In contrast, MVs purified from the Δ*psm*α/*β*Δ*hld* (pTX-*psm*α) or Δ*psm*α/*β*Δ*hld* (pTX-*psmβ*) cultures exhibited no appreciable fluorescence in the presence or absence of xylose. These data demonstrate that only δ-toxin promoted amyloid fibril formation in *S. aureus* planktonic cultures.

Because *S. aureus* extracellular DNA (eDNA) promotes the formation of amyloid fibrils under biofilm–conditions (44), we cultivated strain LAC in the presence or absence of DNase I or RNase. Separate cultures were grown in the presence of proteinase K to digest extracellular proteins, including δ-toxin. MVs from post-exponential cultures were harvested by ultracentrifugation and visualized by TEM. Whereas fibrils were present in MV samples from untreated *S. aureus* cultures or those treated with RNase, the fibrillar content was reduced when the bacteria were cultivated in the presence of DNase or proteinase K (**Fig. 3C).** Likewise, ThT fluorescence of MVs purified from cultures incubated with DNase or proteinase K was markedly reduced compared to that of WT MVs (**Fig. 3D**). Because both enzymes prevented fibril formation during bacterial growth, our results suggest that δ-toxin and eDNA modulate amyloid fibril formation in *S. aureus* planktonic cultures.

### δ-toxin is a major structural component of amyloid fibrils

Because fibrils were associated with MVs purified from WT LAC, Δ*psmα*, and Δ*psmβ*, but not with MVs purified from LACΔ*hld*, we examined whether there were differences in their MV protein profiles as assessed by SDS-PAGE. Untreated MV-associated fibrils did not enter the gel, but samples boiled with SDS and a reducing reagent disaggregated the fibrils prior to sample loading. Four to six major protein bands (~30 to 80 kDa) were observed in MV samples from LAC and its PSM mutants (**Fig. 4A**). Of note, a protein band with a molecular mass <10 kDa was observed in MVs purified from WT LAC, Δ*psmα*, and Δ*psmβ* but not from LACΔ*hld*, suggesting that this band was δ-toxin. Purified MV samples were untreated or digested with DNase, RNase, or proteinase K before analysis by SDS-PAGE. As shown in **Fig. 4B,** DNase or RNase treatment did not markedly alter the protein profile of MV samples. In contrast, the 30 to 80 kDa bands were degraded by proteinase K, whereas the <10 kDa band resisted degradation. The <10 kDa protein band was excised and subjected to LC-MS/MS, and δ-toxin represented >99.9% of its components (**Table S1**). When purified MV preparations were treated for 1 h with proteinase K, DNase I, or RNase before TEM imaging, amyloid fibrils remained in both treated and untreated samples (**Fig. 4C**). Our results indicate that DNase I and proteinase K prevent fibril formation during bacterial growth, but that once mature fibrils are formed, they resist degradation by both enzymes.

**Figure 4.**
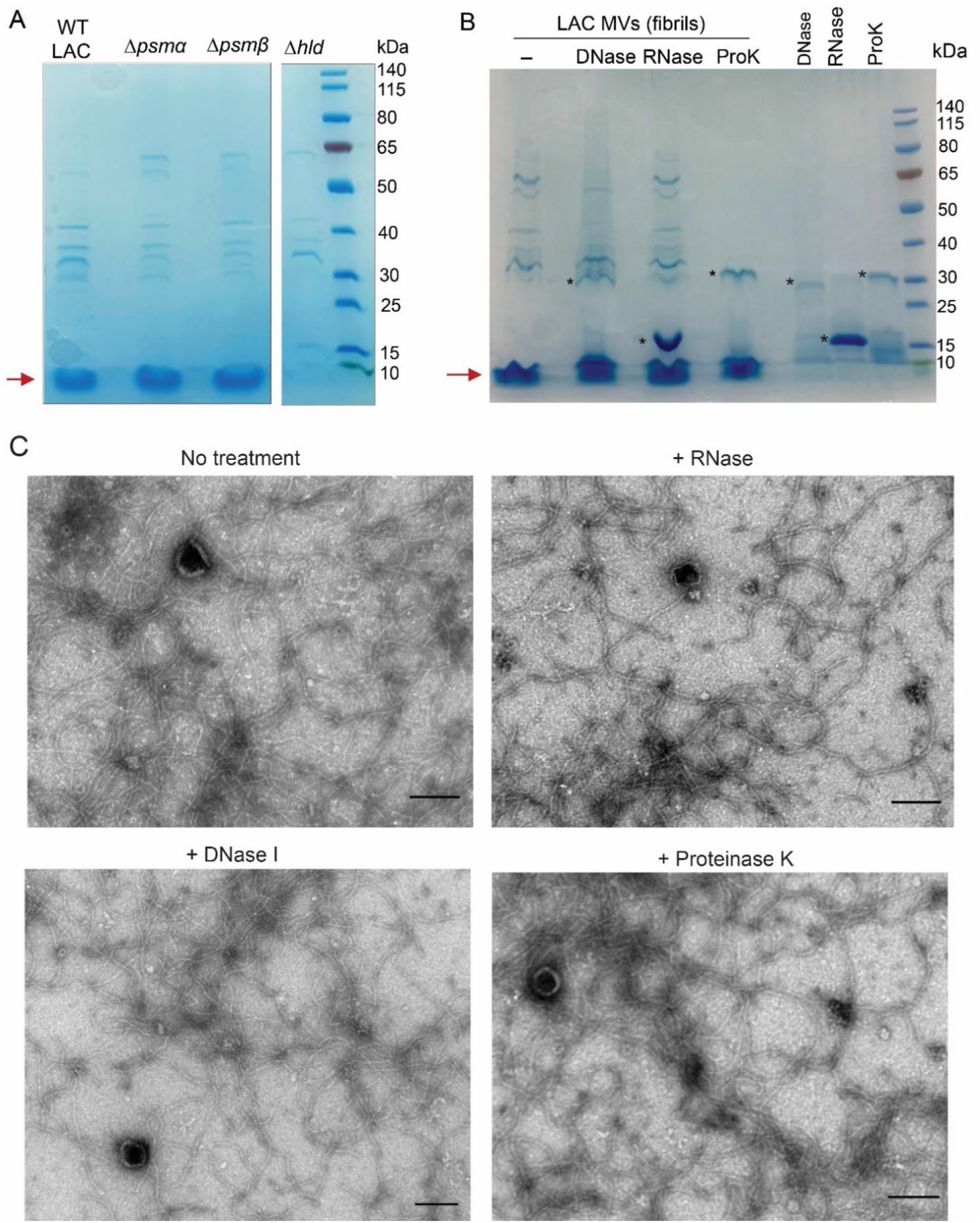
Analysis of enzyme-treated and untreated *S. aureus* MV samples. (A) SDS-PAGE of MVs purified from post-exponential cultures of WT LAC and its PSM mutants or (B) purified LAC MV samples treated with DNase, RNase, or proteinase K (ProK). A band with a molecular weight <10 kDa is indicated by a red arrow. *, indicates major bands associated with enzymes used for MV treatments. (C) Electron micrographs of purified LAC MVs before or after treatment with 200 μg/ml ProK, DNase I, or RNase. Scale bar, 100 nm.

### *S. aureus* PSMα peptides and δ-toxin promote MV production

Mutation of the LAC *psmα* genes significantly reduced MV protein yield (**Fig. 5A**) and particle numbers (**Fig. 5B**) to a greater extent than mutation of the *psmβ* genes. The Δ*hld* mutant showed the lowest MV protein yield and particle numbers, equivalent to that of the triple mutant Δ*psmα/βΔhld*. We extended these findings by complementing the Δ*psmα/βΔhld* mutant with genes encoding either PSMα peptides, PSMβ peptides, or δ-toxin. Gene expression in the complemented mutants was induced with 0, 0.1, 0.3, or 0.5% xylose. As shown in **Figs. 5C and 5D**, the Δ*psmα/βΔhld* mutant carrying either the empty vector pTX or pTX-*psmβ* showed minimal MV yields. In contrast, abundant MV production occurred in the Δ*psmα/βΔhld* (pTX-*psm*α) cultures when the *psmα1-4* genes were induced by 0.1 to 0.5% xylose. MV yields and particle numbers from induced cultures were significantly higher than that of uninduced cultures. Induction of *hld* in Δ*psm*α*/β*Δ*hld* (pTX-*hld*) cultures resulted in xylose inducible, dose-dependent increases in MV production (**Fig. 5C and 5D**). MVs from the Δ*psmα/βΔhld* mutant complemented with pTX-*psmα*, pTX-*psm*β, or pTX-*hld* and induced with 0.5% xylose were visualized by TEM (**Fig. S4**).

**Figure 5.**
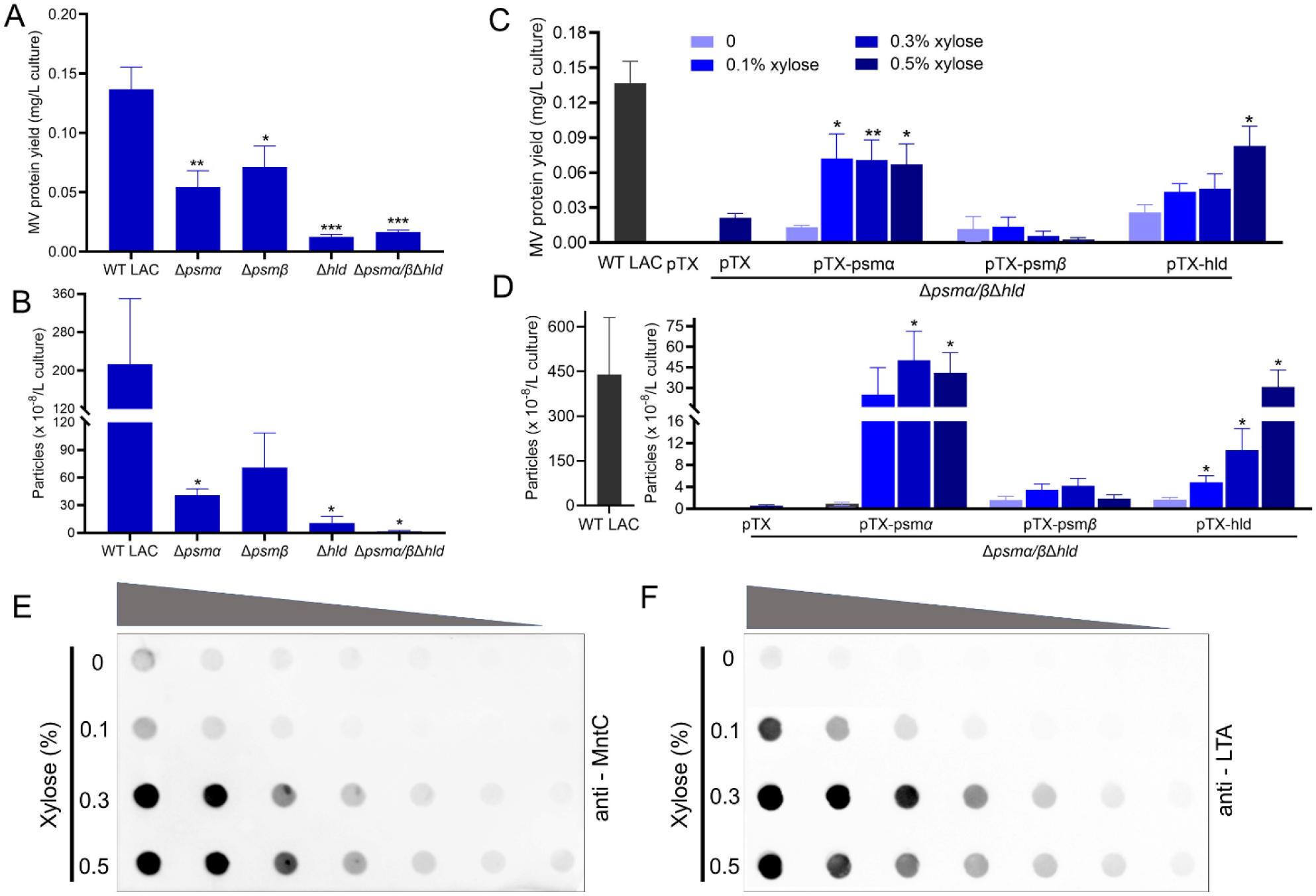
The effect of PSM peptides on the yield of *S. aureus* MVs. Production of MVs from WT LAC and its isogenic mutants lacking *psmα, psmβ*, or *hld* was quantified by (A) total MV protein abundance or (B) MV particle numbers assessed by nanoparticle tracking analysis (n=3). MV production from cultures of the LACΔ*psmα/β*Δ*hld* mutant complemented with *psmα, psmβ*, or *hld* genes induced with indicated concentrations of xylose was quantified by (C) total MV protein yield or (D) MV particle numbers (n=3-5). Two-fold serial dilution of MV samples purified from Δ*psmα/β*Δ*hld* (pTX-*hld*) cultures induced by indicated concentrations of xylose were evaluated by dot blots probed with anti-MntC sera (E) or anti-lipoteichoic acid (LTA) antibodies (F). The dot immunoblot assay was performed at least twice with similar results; a representative blot is shown. MV protein yield and MV particle quantification experiments were expressed as mean + SEM. The data were analyzed using one-way ANOVA with Dunnett’s multiple comparison test; \**P* <0.05, \*\**P* <0.01, \*\*\**P* < 0.001.

As an alternative approach, we utilized dot immunoblots to estimate relative MV concentrations in samples of LAC Δ*psmα/βΔhld* (pTX-*hld*) cultivated in increasing concentrations of xylose. A xylose dose-dependent increase in signal was observed in samples probed with antibodies to the lipoprotein MntC (**Fig. 5E**) or lipoteichoic acid (**Fig. 5F**), antigens abundant in *S. aureus* MVs (16, 26). These findings support our conclusion that induction of δ-toxin by xylose enhances MV production.

### Detection of *S. aureus* MVs in an air pouch infection model

Although many *S. aureus* isolates produce MVs in vitro (12–15, 17, 23, 27), the generation of MVs in vivo during staphylococcal infection is unproven. Visualization of MVs in vivo is difficult because these nanoparticles can only be seen by electron microscopy. Mammalian cells also secrete vesicles (exosomes and microvesicles) (45), and this represents a challenge for the purification of *S. aureus* MVs from body fluids. Synthetic PSM peptides form amyloid fibrils in vitro, but the formation of fibrils in vivo has not yet been documented.

We employed a murine air pouch infection model (**Fig. S5A**) to evaluate in vivo production of MVs. Air pouches were inoculated with live or heat-killed (HK) *S. aureus* LAC at an inoculum of ~10^8^ CFU/mouse. The mice were euthanized 48 h after inoculation, and the pouches were lavaged with 1 ml PBS and cultured quantitatively. Viable bacteria were not recovered from control mice given 10^8^ CFU HK *S. aureus*. Samples from mice that showed bacterial replication in vivo were pooled. After centrifugation of the lavage fluids to remove bacteria and murine cells recruited to the infection site, the supernatant fluids were filter sterilized (0.45 μm). The samples were ultracentrifuged and purified by density gradient ultracentrifugation. MVs were visualized by TEM in samples purified from mice inoculated with *S. aureus* LAC (**Fig. 6A**) but not from mice given HK LAC (**Fig. 6B**), consistent with previous reports that MVs are only generated by live bacteria (46).

**Figure 6.**
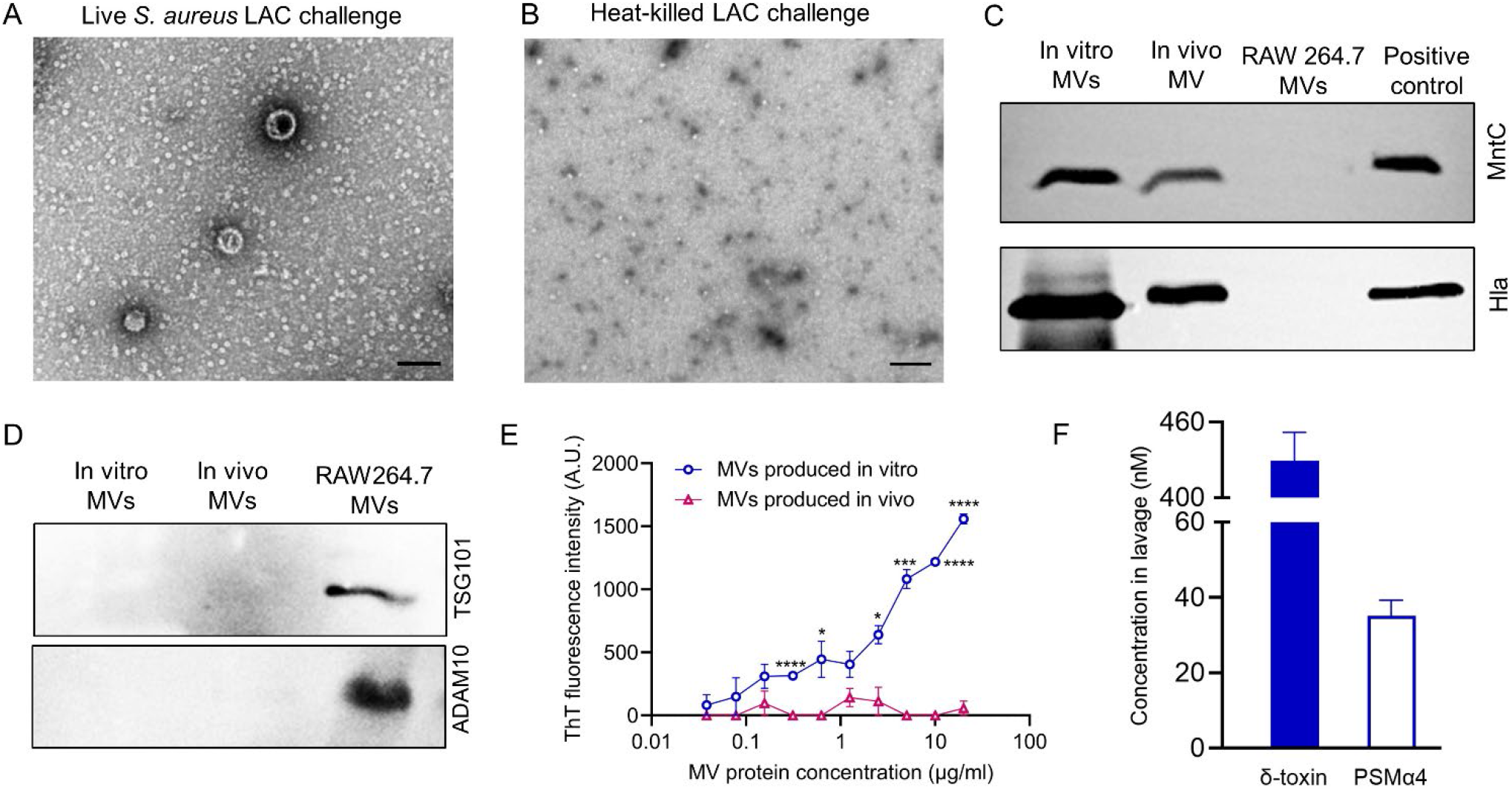
*S. aureus* MVs are generated in vivo in a murine air pouch infection model. Electron micrographs of MV samples purified from pouch lavage fluids of mice infected with viable (A) or heat-killed *S. aureus* LAC (B). Scale bar, 100 nm. (C) Immunoblots show *S. aureus* MV reactivity with antibodies to alpha toxin (Hla) and MntC. (D) Eukaryotic MV markers (TSG101 and ADAM10) only reacted with MVs purified from murine RAW264.7 macrophages. Data from panels A, B, C, and D are representative of at least two independent experiments. (E) ThT fluorescence data were expressed as means ± SEM (n=3) and analyzed by the Student *t*-test. \**P* < 0.05, \*\*\**P* < 0.001, \*\*\*\**P*<0.0001. (F) Concentrations of indicated PSM peptides in the air pouch lavage fluids.

To assess whether MVs recovered from mouse air pouch lavage fluids were microbial in origin, the samples were analyzed by immunoblots. MVs purified from infected mice were reactive with antibodies to *S. aureus* alpha toxin and MntC (**Fig. 6C**), antigens known to be associated with *S. aureus* LAC MVs generated in vitro (16, 26). Antibodies to the eukaryotic vesicle markers ADAM10 (47) and TSG101 (48) were reactive with vesicles purified from culture supernatants of murine RAW 264.7 macrophages (**Fig. 6D**), but not with MVs harvested from air pouch lavage fluids or in vitro *S. aureus* cultures. These data indicate that MVs purified from air pouch infections were *S. aureus* in origin. Of note, amyloid fibrils were not observed in crude MVs directly pelleted from pouch lavage fluids (**Fig. S5B**) or in purified “in vivo” MV samples (**Fig. 6A**). To confirm this observation, MVs harvested either from in vitro cultures or infected air pouches were serially diluted and stained with ThT. MV samples purified from in vitro cultures showed a dose-dependent increase in ThT fluorescence, whereas ThT fluorescence was not measurable in MV samples purified from infected mice (**Fig. 6E**), indicating that PSM amyloids were not formed in this infection model. To determine whether δ-toxin was present in the lavage fluids from infected mice, we analyzed pooled samples by RP-HPLC. Only δ-toxin and PSMα4 were detected in the lavage fluids (**Fig. 6F**), and the δ-toxin concentration in vivo (429 ± 22 nM) was >8-fold higher than that of in vitro post-exponential cultures (51 ± 8 nM).

## Discussion

As a family of small amphipathic, surfactant-like toxins, PSM peptides have multiple biological activities, including cytolysis (28), stimulation of host inflammatory responses, modulation of host innate and adaptive immunity (49), and effects on biofilm maturation (43). PSM monomers, especially PSMα and PSMβ peptides, have been shown to self-assemble into amyloid-like fibrils in vitro, and it has been reported that these amyloid fibrils stabilize in vitro biofilms (34–36). PSMs contribute to in vitro biofilm structuring and detachment, as well as *S. aureus* dissemination from in vivo biofilms (43, 50). Because evidence is lacking that PSM amyloid formation occurs in biofilms formed in vivo, whether this process is biologically relevant remains questionable.

Zhou et al. ammonium sulfate-precipitated PSMs from overnight culture supernatants of the community-acquired *S. aureus* strain MW2, and fibrils composed of δ-toxin were identified in ethanol-extracted precipitates (51). In their studies, formylated δ-toxin formed fibrils, whereas deformylated δ-toxin formed oligomer complexes with PSMα peptides. Only the formylated δ-toxin fibrils bound the amyloid-indicator dye ThT (51). Somerville et al. reported that δ-toxin accumulates in the culture medium in formylated and deformylated forms during the exponential phase of bacterial growth, whereas formylated δ-toxin accumulates during the post-exponential growth phase (52). These observations are consistent with our findings that fibrils were produced in cultures grown to the post-exponential growth phase when formylated δ-toxin levels exceeded those of the other PSMs.

Unlike data generated with synthetic PSMα1, PSMα3, PSMβ1, and PSMβ2 peptides that form amyloid fibrils in vitro (33, 35, 36), our data indicate that amyloid fibril formation relied on dose-dependent *hld* expression in planktonic *S. aureus* LAC cultures. δ-toxin is abundant in culture supernatants of virulent, community-acquired *S. aureus* strains, and for many strains it is the most abundant secreted protein (28, 53). Amyloid fibrils formed in post-exponential cultures of strain LAC bound ThT and were composed primarily of δ-toxin. No fibrils were detected in MVs purified from cultures of hospital-associated *S. aureus* isolates, which are notable for lower levels of PSMs, including δ-toxin (28).

Because bacterial culture supernatants are complex, we postulated that factors other than δ-toxin in the spent culture medium could influence PSM amyloid formation. Schwartz et al. reported that eDNA interacted with PSMα1 to promote amyloid formation in *S. aureus* biofilms, and that an Δ*atl* mutant lacking eDNA did not form fibrils (44). Our data indicate that MV samples purified from Δ*atl* planktonic cultures still contained fibrils (**Fig. S2D**), and thus distinct mechanisms of fibril formation occur in planktonic cultures. When we cultivated *S. aureus* LAC in the presence of DNase I or proteinase K, these enzymes inhibited fibril formation in growing cultures. *S. aureus* MVs are associated with nuclease-susceptible bacterial DNA (54), and thus our findings suggest that eDNA may interact with δ-toxin to promote amyloid fibril formation in *S. aureus* cultures. In contrast, when we treated purified MV preparations with DNase I, RNase, or proteinase K, the amyloid fibrils were resistant to degradation. This result is consistent with previous reports demonstrating that eDNA is protected from DNase-mediated degradation by its interaction with PSM peptides (40), and that amyloid fibrils are resistant to proteinase K degradation (55, 56).

Several studies have reported that PSMα peptides are essential for *S. aureus* MV biogenesis (14, 31). Although δ-toxin proved to be integral to the formation of amyloid fibrils in *S. aureus* cultures, our data indicate that, like PSMα (14), δ-toxin also plays a key role in MV biogenesis. A LACΔ*hld* mutant was equivalent to the triple PSM mutant in terms of reduced MV yield and particle numbers (**Fig. 5A**). WT levels of MVs were restored to the PSM-negative mutant by complementation with either *PSMα1-4* or *hld* (**Fig. 5C**). PSMs are believed to promote MV production by targeting the cytoplasmic membrane and increasing membrane fluidity due to their surfactant-like activity. Indeed, a LACΔ*psm*α/β/*hld* triple mutant showed significantly reduced membrane fluidity compared to the WT strain, and the addition of synthetic PSMα3 or PSMα2 enhanced bacterial membrane fluidity in a dose-dependent fashion (31). An *agr* mutant lacks PSMs due to strict regulation of *psm* expression by AgrA (57), and JE2Δ*agr* showed a ~70% reduction in MV protein yield.

We employed a murine air pouch infection model to determine whether *S. aureus* MVs were generated in vivo and whether “in vivo” MVs might be associated with amyloid fibrils. The air pouch infection model offers the benefit of an in vivo compartment that is easy to sample, and it has been utilized to study inflammation (58), bacterial pathogenesis, host responses to infection (59, 60), and the protective efficacy of multiple vaccine antigens (60). Strain LAC MVs were produced in this infection model, but whether the generation of MVs impacts the pathogenesis of staphylococcal infection remains to be determined.

We did not detect amyloid fibrils associated with MVs harvested from the murine air pouch infection model. Although δ-toxin was the most abundant PSM in lavage fluids, host-derived factors could impede fibril formation in vivo. Najarzadeh et al. reported that human plasma fibrinogen inhibits fibrillation of PSMα1, PSMβ1, and PSMβ2 and induces fibrillation in PSMα3, but its impact on δ-toxin fibrillation was not investigated (61). Serum lipoproteins have been shown to bind to and neutralize the biologic activities of PSMs (62).

In summary, our study investigates the relationship between amyloid formation by PSM peptides and their contribution to the generation of MVs in *S. aureus* cultures. Our work revealed that δ-toxin was responsible for fibril formation in cultures of strain LAC and that, like PSMα, was critical for MV production in vitro. We isolated and purified *S. aureus* MVs from body fluids of animals experimentally infected with *S. aureus*, providing evidence that MVs are produced in vivo. However, amyloid fibrils were not observed in association with MVs purified from infected mice. Our findings provide fundamental insights into the generation of MVs during infection to further our understanding of the contribution of MVs to staphylococcal pathogenesis. The infection model that we used to evaluate MV production in vivo can also be used to investigate MV production by other bacterial pathogens.

## Materials and Methods

### Bacterial strains and growth conditions

*S. aureus* strains (listed in **Table S2**) were cultivated with aeration in tryptic soy broth (Difco) at 37°C. For induction of expression of *psmα, psmβ*, or *hld* in the LACΔ*psmα/βΔhld* mutant, the strains were cultivated at 37°C in Luria broth with 0, 0.1, 0.3, or 0.5% xylose and supplemented with 12.5 μg/ml tetracycline. To complement LACΔ*hld*, pTX-*hld* was transformed into RN4220 by electroporation and then transduced to LACΔ*hld* with φ80α.

### Purification of MVs generated in vitro

*S. aureus* strains were grown at 37°C to the exponential (OD_650nm_ = 0.9; 4 h), post-exponential (OD_650nm_ = 1.2; 4.5 h), or stationary (8 h) phase of growth. Filter-sterilized culture supernatants were concentrated 40-fold by tangential flow filtration with a 100-kDa polyether sulfone membrane system (Centramate, Pall Corp.). The concentrated supernatants were ultracentrifuged at 150,000 × g at 4°C for 3 h to pellet the crude MVs. To remove membrane fragments and protein aggregates, the pellet was gently suspended in PBS and purified by density-gradient ultracentrifugation with 40% to 15% Opti-prep medium. After centrifugation at 140,000 × g for 16 h at 4°C, aliquots of 1 ml gradient fractions were analyzed by SDS-PAGE and silver staining. Fractions with similar protein profiles were pooled, and Optiprep was removed by diafiltration with PBS. Purified MVs were filtered (0.45 μm) and stored at −80°C. MVs were visualized by TEM as described previously (14), and protein concentrations were determined with Bio-Rad protein dye. MV particle enumeration was performed using a Zetaview Nanoparticle Tracking Analyzer (Particle Metrix) with the following settings: camera sensitivity, 85.0; shutter, 70.0; frame rate, 30 frames per second; and temperature, 25°C. Analyses were performed with the in-built Zetaview software 8.02.31.

### ThT fluorescence analysis

To detect amyloid fibrils, 50 μl of PBS containing 0.4 mM ThT (Thermo Fisher) was mixed with an equal volume of MVs or PBS in black 96-well flat bottom plates. After a 30-min incubation at room temperature, the fluorescence of ThT (Ex/Em = 438 nm/495 nm) was measured.

### Quantification of PSM peptides

TCA-precipitated culture supernatants and lyophilized lavage fluid samples were dissolved in 6 M guanidine hydrochloride and analyzed in triplicate by reversed-phase HPLC/MS as described (63), but with a 2.1 x 50 mm C18 column for lavage fluid analysis. PSMs were quantified using Agilent MassHunter software by summing the extracted ion chromatogram (EIC) peak area from two ionized (m/z) species per PSM. PSM concentrations in the samples were determined by calibration with standard curves of synthetic N-formylated PSMs.

### Enzyme treatments

*S. aureus* was cultivated at 37°C to the post-exponential phase of growth in 50 ml tryptic soy broth supplemented with or without 200 μg/ml DNase I, RNase, or proteinase K. Filter-sterilized culture supernatants were ultracentrifuged at 150,000 × g at 4°C for 3 h to pellet the MVs. The samples were resuspended in 100 μl sterile PBS, and the presence of amyloid fibrils was analyzed by TEM and ThT fluorescence. Purified MV samples containing amyloid fibrils were incubated for 1 h at 37°C with or without 200 μg/ml DNase I, RNase, or proteinase K before examination by TEM.

### Dot immunoblot analysis of MV yield

LACΔ*psmα/βΔhld* (pTX-*hld*) cultures were induced with 0, 0.1, 0.3, or 0.5% xylose and cultivated to the post-exponential growth phase. MVs from each culture were purified and concentrated to the same volume. Aliquots (100 μL) of serial twofold dilutions of the MV samples were applied to nitrocellulose membranes using a 96-well Bio-dot apparatus (Bio-Rad). Immunodetection of MntC or LTA was performed as described previously (26).

### Isolation and purification of MVs generated in vivo in the murine air pouch infection model

Animal experiments were carried out in accordance with the recommendations in the PHS Policy on Humane Care and Use of Laboratory Animals. Our animal use protocol (2016N000429) was approved by the Brigham and Women’s Hospital Institutional Animal Care and Use Committee. Air pouches were created by the subcutaneous injection of 3 ml air to the dorsolateral region of mice on days 0 and 3. *S. aureus* strain LAC was cultivated to the logarithmic phase of growth, and the bacteria were harvested and washed to remove MVs generated in vitro. Air pouches were inoculated on day 5 with 0.5 ml of *S. aureus* containing ~10^8^ CFU. After 48 h, the mice were euthanized, and the pouches were lavaged with 1 ml PBS. An aliquot of each air pouch lavage sample was serially diluted and plated to quantify the *S. aureus* CFU/ml. Samples that showed ≥2-fold bacterial growth in vivo were pooled and centrifuged to remove bacteria and host cells. To isolate and purify *S. aureus* MVs generated in vivo, filter sterilized pouch lavage fluids were subjected to ultracentrifugation (150,000 x g) at 4°C for 3 h. The crude MV pellet was gently suspended in PBS and further purified by “top down” OptiPrep density gradient ultracentrifugation, as outlined previously (64). Aliquots of each OptiPrep fraction were subjected to SDS-PAGE and evaluated by immunoblots with polyclonal antibodies to alpha toxin (Hla) (65) (1 μg/ml) or MntC (26). Reactive fractions (6 to 9) were pooled, and the OptiPrep medium was removed by diafiltration with PBS. The samples were negatively stained and examined by TEM.

To validate whether MVs recovered from air pouches were microbial in origin, 5 μg MVs purified from in vitro LAC cultures or from pooled air pouch lavage fluids were subjected to SDS-PAGE and analyzed by immunoblots. Controls included recombinant His-Hla and His-MntC and eukaryotic vesicles purified from culture supernatants of murine RAW 264.7. Eukaryotic MV markers were detected by rabbit anti-TSG101 monoclonal antibody (1 μg/ml; Thermo Fisher; MA1-23296) and rabbit anti-ADAM10 monoclonal antibody (1 μg/ml; Thermo Fisher; MA5-32616).

## Data availability

Data supporting the findings of this manuscript are available from the corresponding author upon reasonable request.

## Acknowledgements

Research reported in this publication was supported by the National Institute of Allergy and Infectious Diseases (NIAID) of the National Institutes of Health under Award R01AI141885 (to J.C.L.) and the NIAID Intramural Research Program project ZIA AI000904 (to M.O.). The content is solely the responsibility of the authors and does not necessarily represent the official views of the National Institutes of Health. We thank William Eagen, Olivia English, and Paul Koffi for their technical assistance. Dr. Victor Torres generously provided the pore-forming toxin mutant *LACΔlukABΔhlgACBΔlukEDΔpvlΔhla*.

## Author contributions

X.W. and J.C.L initiated the project, and X.W., J.C.L, and M.O. designed experiments. X.W., D.U, S.W.D., and D. B. performed the experiments. All authors analyzed the data and wrote the manuscript.

