## Supplement figures and tables for "*Staphylococcus aureus* Delta Toxin Modulates both Extracellular Membrane Vesicle Biogenesis and Amyloid Formation"

### Supplementary Figures

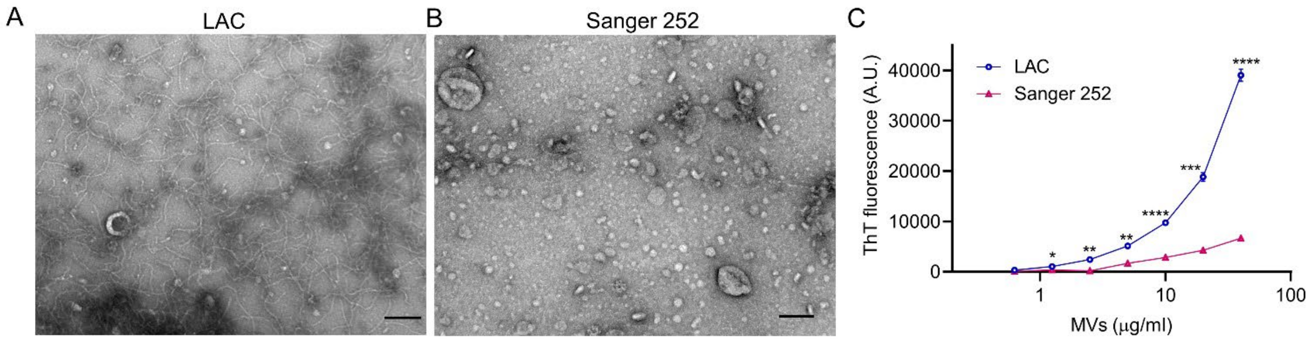


**Figure S1.** Comparison of MV-associated amyloid fibril formation by two *S. aureus* strains. Electron micrograph of crude MVs prepared from post-exponential cultures of (A) *S. aureus* LAC or (B) Sanger 252. Scale bar, 100 nm. (C) ThT fluorescence of MVs purified from LAC and Sanger 252 was expressed as the mean ± SEM (n=3) and analyzed using the Student *t*-test. **P* < 0.05, ***P* < 0.01, ****P* < 0.001, *****P*<0.0001.

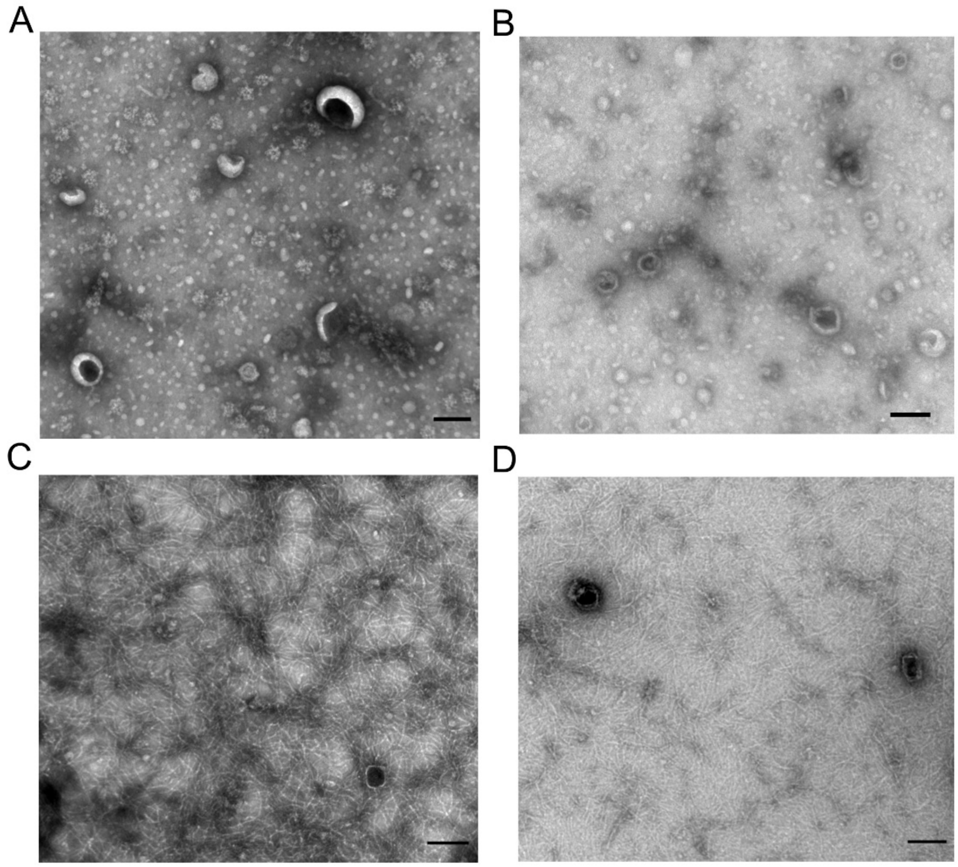


**Figure S2.** Electron micrographs of MV samples purified from *S. aureus* broth cultures. MVs purified from (A) strain LAC cultivated to exponential phase; (B) JE2Δ*agr* cultivated to post-exponential phase

(4.5 h); (C) LACΔ*lukAB*Δ*hlgACB*Δ*lukED*Δ*pvl*Δ*hla* cultivated to post-exponential phase; and (D) JE2Δ*atl* cultivated to post-exponential phase. Scale bars, 100 nm.

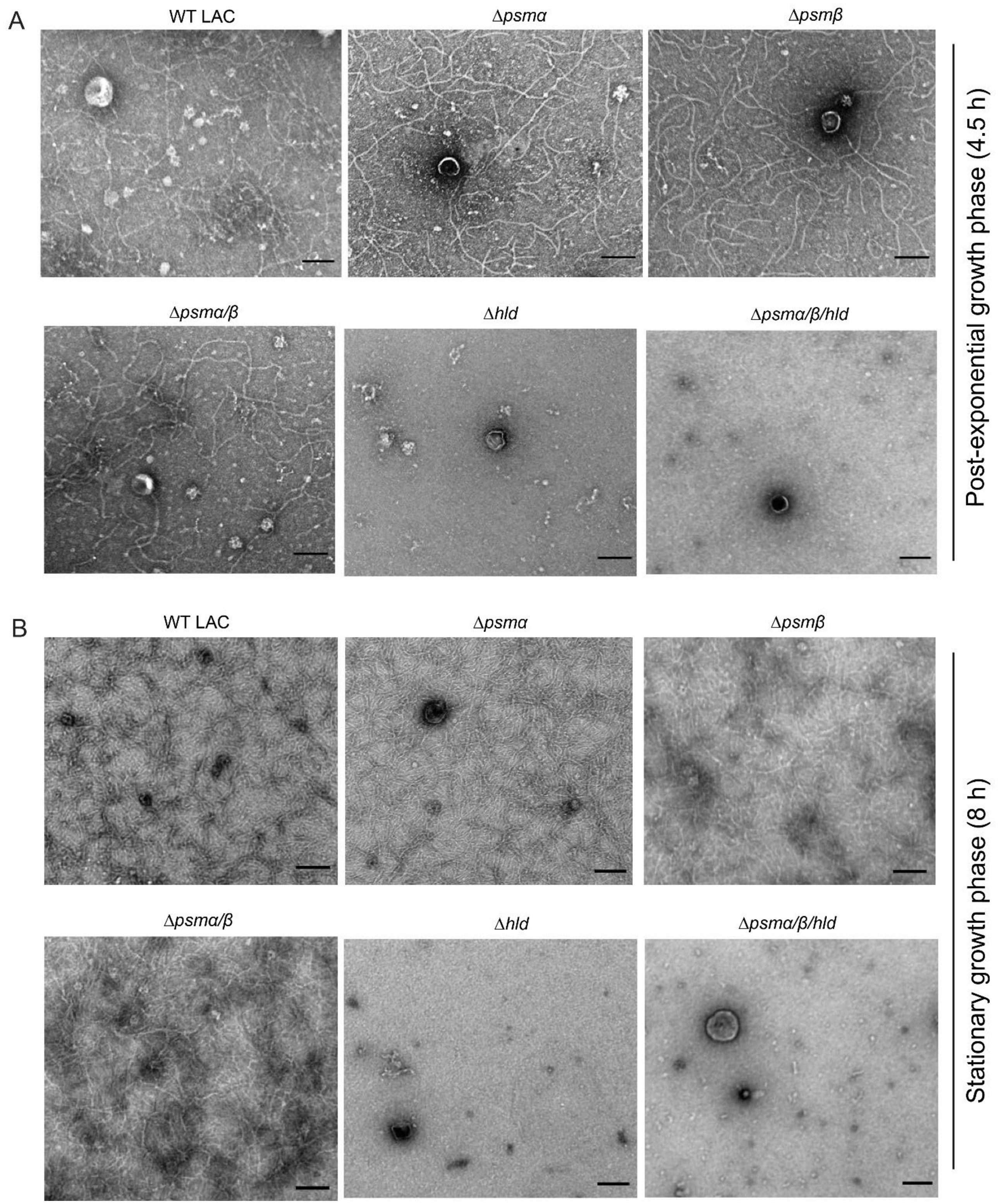

**Figure S3** Electron micrographs of crude MV samples prepared from *S. aureus* LAC and the indicated PSM mutant cultures grown to the post-exponential (A) or stationary growth phase (B). Scale bar, 100 nm.

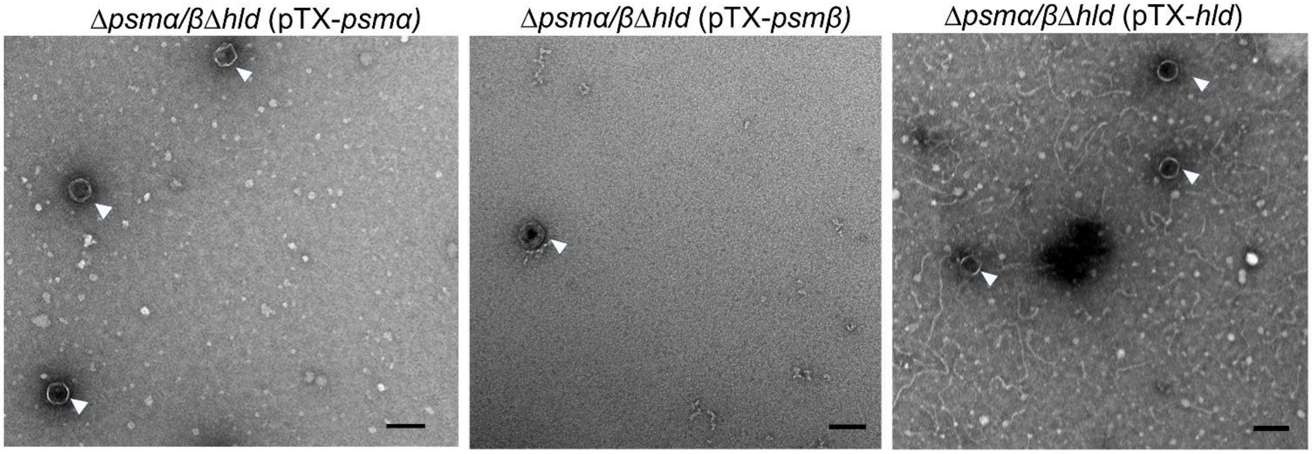


**Figure S4**. Electron micrographs of MV samples purified from cultures of the Δ*psmα/β*Δ*hld* mutant complemented with pTX-*psmα*, pTX-*psmβ*, or pTX-*hld* and induced with 0.5% xylose. MVs in the images are marked with white arrowheads. Scale bar, 100 nm.

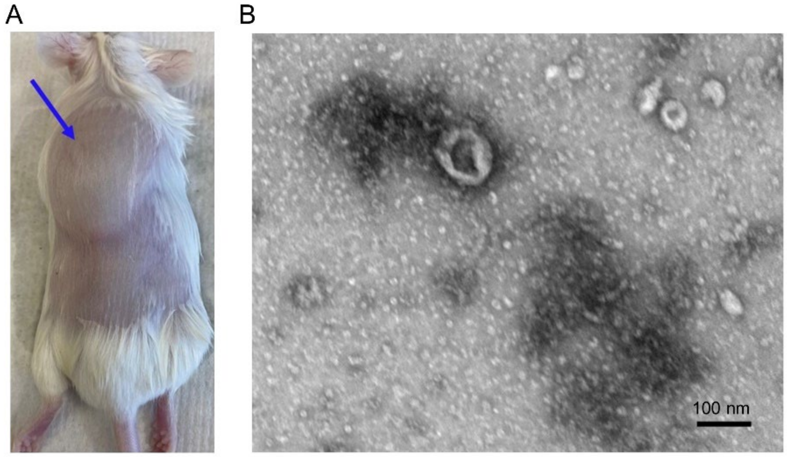

**Figure S5.** The generation of *S. aureus* MVs in an air pouch infection model. (A) The murine air pouch (arrow) prior to bacterial inoculation. (B) Electron micrograph of crude MVs harvested from air pouch lavage fluids of mice challenged with ~$10^8$ CFU viable *S. aureus* LAC. Scale bar, 100 nm.

### Supplementary Tables

**Supplementary Table 1.** Proteins identified in the SDS-PAGE gel band of <10 kDa

| Unique peptides | Total peptides | Annotation | Gene | MW (kDa) | Sum intensity | Intensity (%) | Normalized intensity (%)[1] |
|---|---|---|---|---|---|---|---|
| 2 | 36 | Delta-hemolysin | *hld* | 5.01 | 4.30E+09 | 99.980 | 99.988 |
| 3 | 3 | Hypothetical protein | *sausa300_1904* | 6.56 | 4.60E+05 | 0.011 | 0.008 |
| 1 | 1 | 30S ribosomal protein S17 | *rpsQ* | 10.17 | 2.00E+05 | 0.005 | 0.002 |
| 2 | 2 | Foldase protein | *prsA* | 35.62 | 2.80E+04 | 0.001 | 0.000 |
| 1 | 1 | Putative lipoprotein | *sausa*300_0992 | 23.86 | 5.10E+04 | 0.001 | 0.000 |
| 2 | 2 | Pyruvate dehydrogenase E1 component subunit beta | *pdhB* | 35.22 | 4.50E+04 | 0.001 | 0.000 |
| 1 | 2 | Sensor histidine kinase | *lytS* | 64.99 | 2.60E+04 | 0.001 | 0.000 |
| 1 | 1 | Phenol soluble modulin beta peptide 1 | *psmβ1* | 4.49 | 1.80E+04 | 0.000 | 0.000 |
| 2 | 2 | 2-oxo acid dehydrogenase subunit E2 | *pdhC* | 46.35 | 2.10E+04 | 0.000 | 0.000 |

[1] normalized by molecular weight

**Supplementary Table 2.** *S. aureus* strains used in this study

| Strain | Description[a] | Reference |
|---|---|---|
| JE2 | USA300 LAC cured of three plasmids | (1) |
| JE2Δ*agr* | JE2 *agr*::*tetM*, Tc$^r$ | (2) |
| JE2Δ*atl* | JE2 *atl*::*bursa aurealis*, Em$^r$ | (1) |
| LAC | CA-MRSA, USA300 | (3) |
| LACΔ*psmα* | *psmα* deletion mutant | (4) |
| LACΔ*psmβ* | *psmβ* deletion mutant | (4) |
| LACΔ*psmα/β* | *psmα/psmβ* double mutant | (5) |
| LACΔ*hld* | *hld* mutant; hld start codon changed from ATG to ATT | (4) |
| LACΔ*psmα/β*Δ*hld* | *psmα/psmβ/hld* triple mutant | (5) |
| LACΔ*hld* (pTX-*hld*) | *hld* gene is expressed with inducible vector pTX Tc$^r$ | This study |
| LACΔ*psmα/β*Δ*hld* (pTX-*psmα1-4*) | *psmα1-4* genes are expressed with inducible vector pTX, Tc$^r$ | (6) |
| LACΔ*psmα/β*Δ*hld* (pTX-*psmβ1-2*) | *psmβ1-2* genes are expressed with inducible vector pTX, Tc$^r$ | (6) |
| LACΔ*psmα/β*Δ*hld* (pTX-*hld*) | *hld* gene is expressed with inducible vector pTX, Tc$^r$ | (6) |
| LACΔ*lukAB*Δ*hlgACB*Δ*lukED*Δ*pvl*Δ*hla* | Pore-forming toxin mutant lacking *hla* and all leukocidin genes | (7) |
| MN8 | MSSA, USA200, ST30 | (8) |
| N315 | HA-MRSA, ST5 | (9) |
| Sanger 252 | HA-MRSA, ST36 | (10) |
| MW2 | CA-MRSA, USA400, ST1 | (11) |
| NRS483 | CA-MRSA, USA1000, ST59 | BEI Resources |
| RN4220 | Restriction-deficient mutant of *S. aureus* 8325-4 | (12) |

[a]Tc, tetracycline; CA-MRSA, community-acquired methicillin resistant *Staphylococcus aureus*; HA-MRSA, hospital-acquired methicillin resistant *Staphylococcus aureus*.
